# Phylogenomics and phylodynamics of Zika Virus from Asia

**DOI:** 10.1101/2023.11.16.567488

**Authors:** Kamil Baig, Shazia Farah, Ashwin Atkulwar

## Abstract

The current study analyzes the Zika virus from Asia from a phylogenomic and phylodynamic perspective. To perform this study, 68 genomes from Asia were analyzed. A similar dataset was used to perform phylogenomics using Median Joining Network (MJN). MJN reconfirms African origin of Zika virus along with few notable instances of common origin of infection in some Asian countries. We investigated population dynamics using SkyGrid, Exponential Growth, and Constant Coalescence models. According to our phylodynamic analysis, effective population sizes throughout Asia reached higher peaks during the outbreak of 2016-2017. Zika virus population size increased exponentially during 2016-2017. From 1966 to 2021, there was a high and low peak in effective population size except for the outbreak of 2016-2017. To counter the outbreak of Zika Virus in Asian Countries in future, we recommend continuous genomic surveillance.

## 1. Introduction

In order to monitor endemic and emerging infectious diseases from a genomic perspective requires molecular epidemiological studies of various viral variants. In studies of the molecular epidemiology of various viruses, the molecular mutations are mostly found to be linked to the various viral phenotypic traits related to infectiousness, and the pathogenicity. The documentation of these emerging mutations is crucial for the implementation of public health strategies to minimize outbreak effects. A number of pioneering studies were conducted to understand the molecular epidemiology, the phylogenomics, and the phylodynamics of a number of human-associated viruses including human immunodeficiency virus HIV-1 [1], hepatitis C virus (HCV) [2], Zika virus (ZIKV) [3], and severe acute respiratory syndrome coronavirus 2 (SARS-CoV-2) [4]. Zika virus is a member of the Flaviviridae virus family, first isolated from a rhesus monkey in Uganda’s Zika forest in 1947 [5, 6, and 7]. Until 2007, the Zika virus was mostly confined to equatorial belt from Africa to Asia, but after 2007, it spread across the Pacific Ocean to America, causing epidemics in 2015 and 2016 [8]. In 2016, the World Health Organization (WHO), declared the outbreak of Zika Virus an international health emergency (http://www.who.int/en/). The Zika Virus is exclusively transmitted by Andes mosquitoes but monkeys can also act as vectors and humans are also act as occasional host. Zika Virus Infection symptoms range from asymptomatic illness to severe conditions, called as Zika fever which causes headaches, rash, malaise chills etc. [9]. Studies suggest that the Zika Virus Infection may also cause neurological defects such as Guillain-Barre syndrome [10], and microcephaly [11, 12]. In addition to zoonotic and mother-to-child transmission, the Zika virus can also be transmitted sexually [13, 14]. Zika virus consists of single-stranded RNA genome that contains 10794 kilo bases and two flanking noncoding regions. A major virion surface protein in Zika genome is the E protein, which facilitates membrane fusion and binding at various stages of the viral cycle [15]. This study examines the phylogenomic and phylodynamic of Zika virus isolates from Asia.

### 2. Material and methods

From the NCBI Virus database 68 whole-genome sequences from Asia were retrieved (Supplementary Table 1). For phylogenetic network analysis a reference genome from Uganda, Africa was also added to the Asian database. For downstream analysis, all genomes were aligned using Clustal Omega [16]. Phylogenomic was performed using Median Joining Network construction by method as implemented in the program NETWORK [17] (Supplementary Table 1). BEAST v1.10.4, software package was used to investigate the evolutionary dynamics of the Zika Virus. To reconstruct evolutionary dynamics, the HKY nucleotide substitution model with a gamma category count of 4 and a strict clock, with a mean clock rate of 1.0 was used. A Bayesian SkyGrid, exponential growth, and constant coalescence tree priors was used to investigate trends in Zika Virus distribution in Asian populations. In order to draw plots, Tracer v1.7.0 was used to read logs and tree files. Tracer v1.7.0 was also used to determine the ESS support for priors and various parameters [18]. As a burn-in step, the first 10% trees were discarded using the MCMC chain length of 10 million steps.

### 3. Result and discussion

The median-joining network (MJN) constructed from 67 haplotypes out of 69 genomes is characterized by the presence of greater number of median vectors (Figure 1). The Median vectors indicate either an unsampled genomes or an extinct virus [19]. It is noteworthy that none of these 69 genomes across Asia (including a reference genome from Uganda Accession No. NC_012532) shared haplotypes to form haplogroups. Based on the absence of haplogroups, it appears that the virus is etiologically different or evolved locally. Compared to the closest Asian genomes, the Ugandan genome differed by more than ∼1000 mutations confirming its different geographical and ancestral position.

**Figure 1.**
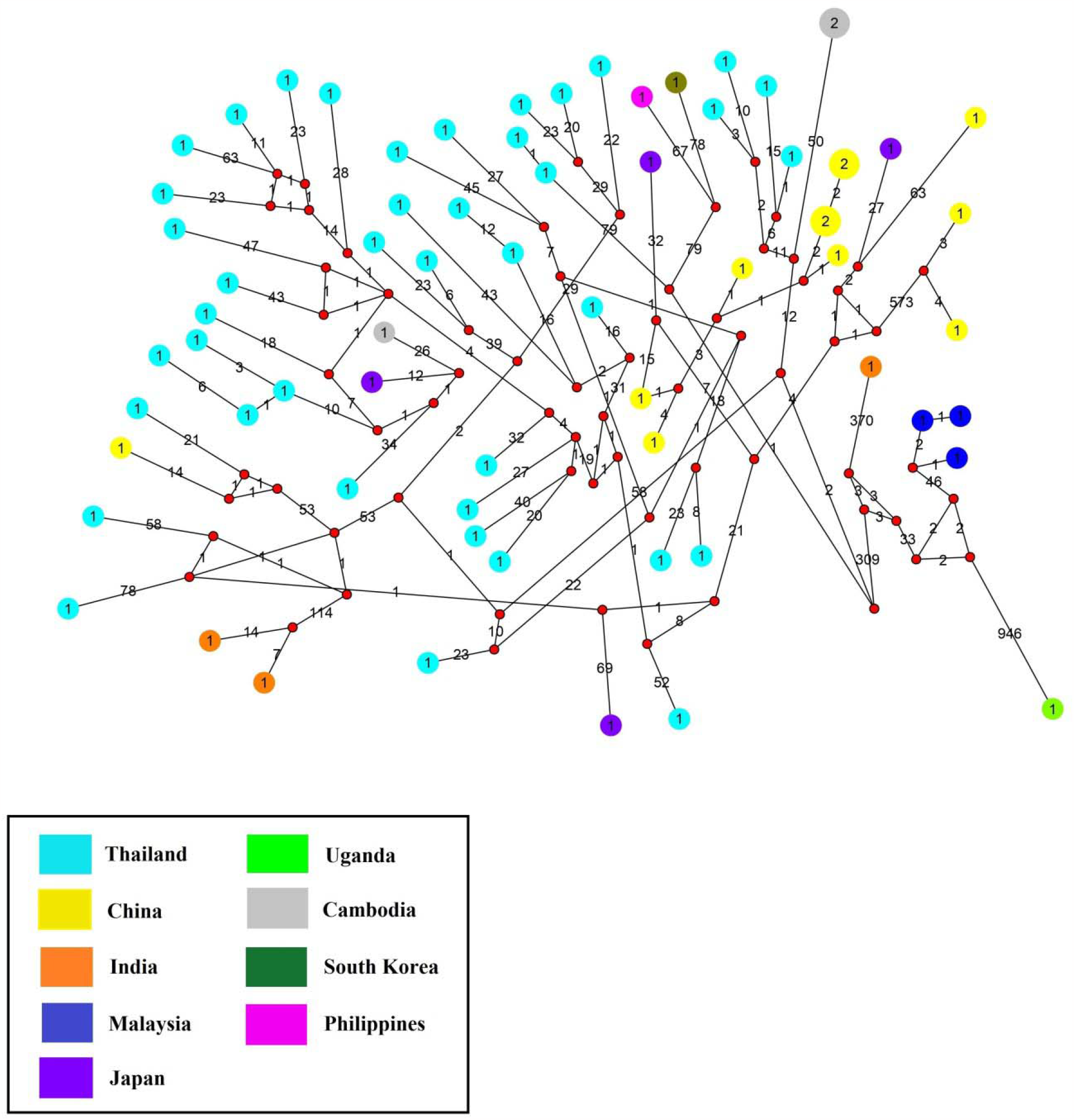
Median Joining Network (MJN) constructed out of 64 genomes of Zika viruses from Asia. The countries are represented by different color circles. The size of the circles corresponds to the number of shared haplotypes. Links with black numbers exhibits number of mutated nucleotides. The solid red dots represent median vectors. Note the highest number of mutated nucleotides in link connecting Uganda genome to the Asiatic Zika genomes. Also the MJN show many median vectors.

China and Cambodia have only two genomes with more than frequency each, while the rest have only one. Few haplotypes from Malaysia, South Korea, Philippines, China, Cambodia and Japan have evolved through common source of origin manifested by single median vector connecting them.

The population dynamics of the Zika virus inferred from the whole genome sequences deposited from Asian countries (between 1966 and 2021) was plotted using a Bayesian Skyline Plot (BSP) method shows irregularities in the Zika virus’s population dynamics (Figure 2). Based on the SkyGrid model, phylodynamic analysis revealed a sigmoidal distribution over a long time scale (from 1966 to 2021), with a relatively constant effective population size observed from 1970 to 1990 (Figure 2 (a). Since 1985 the constant effective population size has decreased and continued to decline until 2000, than increased exponentially until 2016. In addition, we found the highest peak of Zika cases throughout Asia in 2016-2017, which declined again in 2018. In a similar period, there have been outbreaks of Zika and sporadic autochthonous transmission events in countries like Bangladesh, Cambodia, India, Indonesia, Laos, Malaysia, Maldives, Myanmar, Philippines, Singapore, Thailand, and Vietnam [20, 21, 22 and 23]. During the period August 27 through November 30, 2016, 455 cases were reported in Singapore (Singapore Zika Study 2017), while in Thailand ∼700 cases were reported the highest in Asia, and there were more than 200 cases reported in Vietnam in 2016 [20]. The exponential growth model, however, indicated that the effective population size increased exponentially from 1966 to 2021 (Figure 2 (b). Constant coalescence model shows more or less the same type of population dynamics as SkyGrid model (Figure 2 (c)). Out study confirms previous findings and also provides new insights into the phylodynamics and phylogenomics of Zika virus despite the limited number and over or under-representation of genomes from Asia. Developing strategies to mitigate Zika outbreaks in Asia would be made possible by widespread sequencing and analyses of genomes from Asia.

**Figure 2.**
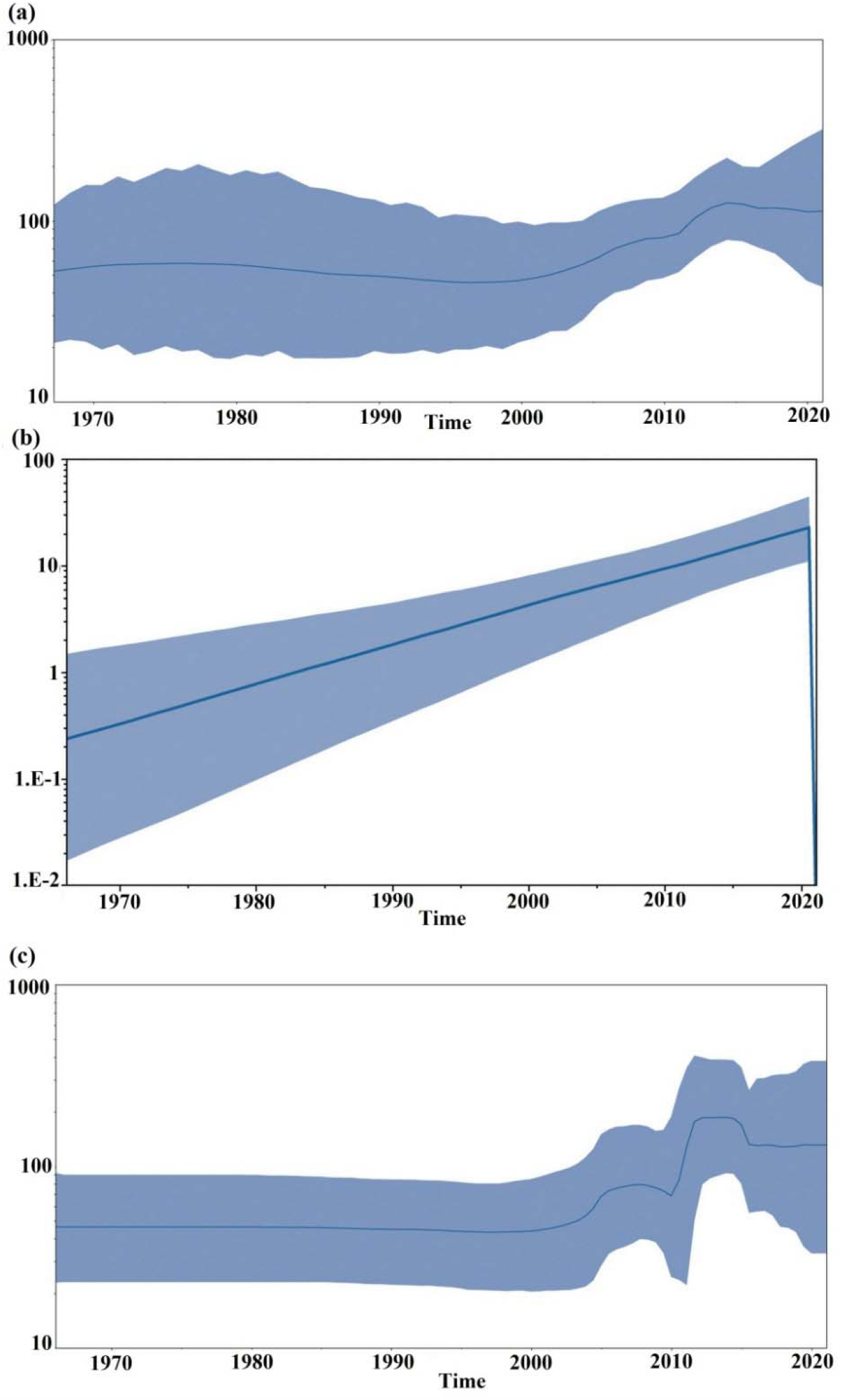
Bayesian skyline plots (a), (b) & (c) constructed using 68 Asian Zika genomes by SkyGrid, exponential growth and coalescence Bayesian skyline tree priors implemented. The Y-axis of the graph denotes effective population size (*N*_*e*_) while the X-axis indicates timeline in years.

## Authors’ contributions

KB performed the analyses, SF performed analyses and wrote the manuscript, AA conceived the idea and wrote the manuscript.

## Acknowledgments

The authors and submitting laboratories of the genomes from GISAID are gratefully acknowledged.

## Reference

1. Theys K, Lemey P, Vandamme AM, Baele G. Advances in Visualization Tools for Phylogenomic and Phylodynamic Studies of Viral Diseases. Front. Public Health. 2019. Infectious Diseases – Surveillance, Prevention and Treatment Volume 7. DOI: 10.3389/fpubh.2019.0020

2. Paraschiv S, Banica L, Nicolae I, Niculescu I, Abagiu A, Jipa R, Pineda-Peña AC, Pingarilho M, Neaga E, Theys K, Libin P, Otelea D, Abecasis A. Epidemic dispersion of HIV and HCV in a population of co-infected Romanian injecting drug users. PLoS One. 2017; 12(10):e0185866. doi: 10.1371/journal.pone.0185866.

3. Aubry F, Dabo S, Manet C, Filipović I, Rose NH, Miot EF, Martynow D, Baidaliuk A, Merkling SH, Dickson LB, Crist AB. Enhanced Zika virus susceptibility of globally invasive Aedes aegypti populations. Science. 2020; 370(6519):991–6.

4. Rambaut A, Holmes EC, O’Toole Á, Hill V, McCrone JT, Ruis C, du Plessis L, Pybus OG. A dynamic nomenclature proposal for SARS-CoV-2 lineages to assist genomic epidemiology. Nature microbiology. 2020; 5(11):1403–7.

5. Malone RW, Homan J, Callahan MV, Glasspool-Malone J, Damodaran L, Schneider AD, Zimler R, Talton J, Cobb RR, Ruzic I, Smith-Gagen J. Zika virus: medical countermeasure development challenges. PLoS neglected tropical diseases. 2016; 10(3):e0004530. doi:10.1371/journal.pntd.0004530

6. Henry R. Zika virus. Emerg Infect Dis. 2014; 20(6):1090. doi: 10.3201/eid2006.et2006. PMID: 24983096; PMCID: PMC4036762.

7. Sikka V, Chattu VK, Popli RK, Galwankar SC, Kelkar D, Sawicki SG, Stawicki SP, Papadimos TJ. The Emergence of Zika Virus as a Global Health Security Threat: A Review and a Consensus Statement of the INDUSEM Joint working Group (JWG). J Glob Infect Dis. 2016; 8(1):3–15. doi: 10.4103/0974-777X.176140.

8. Mehrjardi ZM, Poretti A, Huisman TA, Werner H, Keshavarz E, Araujo Júnior E. Neuroimaging findings of congenital Zika virus infection: a pictorial essay. Jpn J Radiol. 2017; 35(3):89–94. doi: 10.1007/s11604-016-0609-4.

9. Hayes EB. Zika virus outside Africa. Emerg Infect Dis. 2009; 15 (9):1347–50. doi: 10.3201/eid1509.090442.

10. Oehler E, Watrin L, Larre P, Leparc-Goffart I, Lastere S, Valour F, Baudouin L, Mallet H, Musso D, Ghawche F. Zika virus infection complicated by Guillain-Barre syndrome--case report, French Polynesia, December 2013. Euro Surveill. 2014;19(9):20720. doi: 10.2807/1560-7917.es2014.19.9.20720.

11. Fauci AS, Morens DM. Zika Virus in the Americas--Yet Another Arbovirus Threat. N Engl J Med. 2016; 374(7):601–4. doi: 10.1056/NEJMp1600297.

12. Ventura CV, Maia M, Bravo-Filho V, Góis AL, Belfort R Jr. Zika virus in Brazil and macular atrophy in a child with microcephaly. Lancet. 2016; 387(10015):228. doi: 10.1016/S0140-6736(16)00006-4.

13. Musso D, Roche C, Robin E, Nhan T, Teissier A, Cao-Lormeau VM. Potential sexual transmission of Zika virus. Emerg Infect Dis. 2015 Feb;21(2):359–61. doi: 10.3201/eid2102.141363. Erratum in: Emerg Infect Dis. 2015; 21(3):552.

14. Foy BD, Kobylinski KC, Chilson Foy JL, Blitvich BJ, Travassos da Rosa A, Haddow AD, Lanciotti RS, Tesh RB. Probable non-vector-borne transmission of Zika virus, Colorado, USA. Emerg Infect Dis. 2011; 17(5):880–2. doi: 10.3201/eid1705.101939.

15. Lindenbach BD, Rice CM. Molecular biology of flaviviruses. Adv Virus Res. 2003; 59:23–61. doi: 10.1016/s0065-3527(03)59002-9.

16. Sievers F, Wilm A, Dineen D, Gibson TJ, Karplus K, Li W, Lopez R, McWilliam H, Remmert M, Söding J, Thompson JD, Higgins DG. Fast, scalable generation of high-quality protein multiple sequence alignments using Clustal Omega. Mol Syst Biol. 2011;7:539. doi: 10.1038/msb.2011.75.

17. Bandelt HJ, Forster P, Röhl A. Median-joining networks for inferring intraspecific phylogenies. Mol Biol Evol. 1999;16(1):37–48. doi: 10.1093/oxfordjournals.molbev.a026036.

18. Rambaut A, Drummond AJ, Xie D, Baele G, Suchard MA. Posterior Summarization in Bayesian Phylogenetics Using Tracer 1.7. Syst Biol. 2018; 67(5):901–904. doi: 10.1093/sysbio/syy032.

19. Atkulwar, A. Rehman, A. Imaan, Y. Baig, M. Analyses of Omicron Genomes from India Reveal BA.2 As a More Transmissible Variant. European Journal of Biological Research. 2023; 13, 10–17.

20. Lim SK, Lim JK, Yoon IK. An Update on Zika Virus in Asia. Infect Chemother. 2017;49(2):91–100. doi: 10.3947/ic.2017.49.2.91.

21. Zanluca C, Melo VC, Mosimann AL, Santos GI, Santos CN, Luz K. First report of autochthonous transmission of Zika virus in Brazil. Mem Inst Oswaldo Cruz. 2015;110 (4):569–72. doi: 10.1590/0074-02760150192.

22. Hu T, Li J, Carr MJ, Duchêne S, Shi W. The Asian Lineage of Zika Virus: Transmission and Evolution in Asia and the Americas. Virol Sin. 2019;34(1):1–8. doi: 10.1007/s12250-018-0078-2.

23. Luo XS, Imai N, Dorigatti I. Quantifying the risk of Zika virus spread in Asia during the 2015-16 epidemic in Latin America and the Caribbean: A modeling study. Travel Med Infect Dis. 2020; 33:101562. doi: 10.1016/j.tmaid.2020.101562.

