## Supplementary Table 1 for "Phylogenomics and phylodynamics of Zika Virus from Asia"

**Supplementary Table S1. Detail information about genomic sequences of Zika Virus, retrieved from the NCBI Virus database, used in this research work.**

| **Sr. No** | **Accession No.** | **Submitter** | **Length** | **Collection Date** | **Geo Location** | **Host** |
| --- | --- | --- | --- | --- | --- | --- |
| 1 | OQ661912 | Klungthong,C., Chinnawirotpisan,P., Manasatienkij,W., Poolpanichupatam,Y., Hussem,K., Lohachanakul,J., Buddhari,D., Suntarattiwong,P., Watanaveeradej,V., Suwanpakdee,D., Chuenchitra,T., Chusri,S., Buathong,R., Fernandez,S., Macareo,L.R | 10699 | ######## | Thailand | Homo sapiens |
| 2 | OQ661913 | Klungthong,C., Chinnawirotpisan,P., Manasatienkij,W., Poolpanichupatam,Y., Hussem,K., Lohachanakul,J., Buddhari,D., Suntarattiwong,P., Watanaveeradej,V., Suwanpakdee,D., Chuenchitra,T., Chusri,S., Buathong,R., Fernandez,S., Macareo,L.R | 10699 | ######## | Thailand | Homo sapiens |
| 3 | OQ661914 | Klungthong,C., Chinnawirotpisan,P., Manasatienkij,W., Poolpanichupatam,Y., Hussem,K., Lohachanakul,J., Buddhari,D., Suntarattiwong,P., Watanaveeradej,V., Suwanpakdee,D., Chuenchitra,T., Chusri,S., Buathong,R., Fernandez,S., Macareo,L.R | 10699 | 7/4/2011 | Thailand | Homo sapiens |
| 4 | OQ661915 | Klungthong,C., Chinnawirotpisan,P., Manasatienkij,W., Poolpanichupatam,Y., Hussem,K., Lohachanakul,J., Buddhari,D., Suntarattiwong,P., Watanaveeradej,V., Suwanpakdee,D., Chuenchitra,T., Chusri,S., Buathong,R., Fernandez,S., Macareo,L.R | 10699 | ######## | Thailand | Homo sapiens |
| 5 | OQ661916 | Klungthong,C., Chinnawirotpisan,P., Manasatienkij,W., Poolpanichupatam,Y., Hussem,K., Lohachanakul,J., Buddhari,D., Suntarattiwong,P., Watanaveeradej,V., Suwanpakdee,D., Chuenchitra,T., Chusri,S., Buathong,R., Fernandez,S., Macareo,L.R | 10699 | ######## | Thailand | Homo sapiens |
| 6 | OQ661917 | Klungthong,C., Chinnawirotpisan,P., Manasatienkij,W., Poolpanichupatam,Y., Hussem,K., Lohachanakul,J., Buddhari,D., Suntarattiwong,P., Watanaveeradej,V., Suwanpakdee,D., Chuenchitra,T., Chusri,S., Buathong,R., Fernandez,S., Macareo,L.R | 10699 | ######## | Thailand | Homo sapiens |
| 7 | OQ661918 | Klungthong,C., Chinnawirotpisan,P., Manasatienkij,W., Poolpanichupatam,Y., Hussem,K., Lohachanakul,J., Buddhari,D., Suntarattiwong,P., Watanaveeradej,V., Suwanpakdee,D., Chuenchitra,T., Chusri,S., Buathong,R., Fernandez,S., Macareo,L.R | 10699 | 2018 | Thailand | Homo sapiens |
| 8 | OQ661919 | Klungthong,C., Chinnawirotpisan,P., Manasatienkij,W., Poolpanichupatam,Y., Hussem,K., Lohachanakul,J., Buddhari,D., Suntarattiwong,P., Watanaveeradej,V., Suwanpakdee,D., Chuenchitra,T., Chusri,S., Buathong,R., Fernandez,S., Macareo,L.R | 10699 | ######## | Thailand | Homo sapiens |
| 9 | OM964565 | Khongwichit,S., Chuchaona,W., Thongmee,T., Vongpunsawad,S., Poovorawan,Y | 10608 | ######## | Thailand | Homo sapiens |
| 10 | OM964566 | Khongwichit,S., Chuchaona,W., Thongmee,T., Vongpunsawad,S., Poovorawan,Y | 10608 | ######## | Thailand | Homo sapiens |
| 11 | OM964567 | Khongwichit,S., Chuchaona,W., Thongmee,T., Vongpunsawad,S., Poovorawan,Y. | 10608 | ######## | Thailand | Homo sapiens |
| 12 | OM964568 | Khongwichit,S., Chuchaona,W., Thongmee,T., Vongpunsawad,S., Poovorawan,Y. | 10608 | ######## | Thailand | Homo sapiens |
| 13 | OP281680 | Yoksan,S., Rodpai,E., Rajkam,S., Punyahathaikul,S., Wanlayaporn,D., Masrinoul,P. | 10807 | ######## | Thailand | Homo sapiens |
| 14 | OM666891 | Yadav,P.D. | 10749 | ######## | India | Homo sapiens |
| 15 | OM666892 | Yadav,P.D. | 10753 | ######## | India | Homo sapiens |
| 16 | OK054351 | Yadav,P.D. | 10748 | ######## | India: Pune City, Maharashtra State | Homo sapiens |
| 17 | MT377491 | Dangsagul,W., Klangkratok,C., Changsom,D., Chanmanee,T., Tawatsin,A., Sangkijporn,S., Puthavathana,P | 10663 | ######## | Thailand: Nakorn Ratchasrima | Homo sapiens |
| 18 | MT377492 | Dangsagul,W., Chanmanee,T., Phatihattakorn,C., Sangsiriwut,K., Tawatsin,A., Sangkijporn,S., Auewarakul,P., Puthavathana,P | 10619 | ######## | Thailand: Bangkok | Homo sapiens |
| 19 | MT377493 | Dangsagul,W., Chanmanee,T., Putchakarn,S., Sirichote,P., Chaisomboonpan,S., Sangsiriwut,K., Tawatsin,A., Sangkijporn,S., Puthavathana,P | 10613 | ######## | Thailand: Samut Sakorn | Homo sapiens |
| 20 | MT377494 | Dangsagul,W., Chanmanee,T., Putchakarn,S., Sirichote,P., Chaisomboonpan,S., Sangsiriwut,K., Tawatsin,A., Sangkijporn,S., Puthavathana,P. | 10736 | ######## | Thailand: Samut Sakorn | Homo sapiens |
| 21 | MT377495 | Dangsagul,W., Chanmanee,T., Putchakarn,S., Sirichote,P., Chaisomboonpan,S., Sangsiriwut,K., Tawatsin,A., Sangkijporn,S., Puthavathana,P. | 10606 | ######## | Thailand: Samut Sakorn | Homo sapiens |
| 22 | MT377496 | Dangsagul,W., Pairoj,T., Chanmanee,T., Sangsiriwut,K., Tawatsin,A., Sangkijporn,S., Puthavathana,P. | 10602 | ######## | Thailand: Chonburi | Homo sapiens |
| 23 | MT377497 | Dangsagul,W., Pairoj,T., Chanmanee,T., Sangsiriwut,K., Tawatsin,A., Sangkijporn,S., Puthavathana,P. | 10602 | ######## | Thailand: Rayong | Homo sapiens |
| 24 | MT377498 | Dangsagul,W., Klangkratok,C., Chanmanee,T., Sangsiriwut,K., Ruchusatsawat,K., Tawatsin,A., Sangkijporn,S., Puthavathana,P. | 10610 | ######## | Thailand: Bueng Kan | Homo sapiens |
| 25 | MT377499 | Dangsagul,W., Klangkratok,C., Chanmanee,T., Sangsiriwut,K., Ruchusatsawat,K., Tawatsin,A., Sangkijporn,S., Puthavathana,P. | 10585 | ######## | Thailand: Nakhon Pathom | Homo sapiens |
| 26 | MT377500 | Dangsagul,W., Klangkratok,C., Chanmanee,T., Sangsiriwut,K., Ruchusatsawat,K., Tawatsin,A., Sangkijporn,S., Puthavathana,P. | 10619 | ######## | Thailand: Phetchaburi | Homo sapiens |
| 27 | MT377501 | Dangsagul,W., Klangkratok,C., Chanmanee,T., Sangsiriwut,K., Ruchusatsawat,K., Tawatsin,A., Sangkijporn,S., Puthavathana,P. | 10591 | ######## | Thailand: Bangkok | Homo sapiens |
| 28 | MT377502 | Dangsagul,W., Changsom,D., Sangsiriwut,K., Chokephaibulkit,K., Puthavathana,P., Yoksan,S. | 10770 | ######## | Thailand | Homo sapiens |
| 29 | MT377503 | Dangsagul,W., Changsom,D., Sangsiriwut,K., Chokephaibulkit,K., Puthavathana,P., Yoksan,S. | 10607 | ######## | Thailand | Homo sapiens |
| 30 | MT377504 | Dangsagul,W., Changsom,D., Sangsiriwut,K., Chokephaibulkit,K., Puthavathana,P., Yoksan,S. | 10603 | 9/6/2006 | Thailand | Homo sapiens |
| 31 | MZ008356 | Yang,F. | 10686 | 2019-12 | Cambodia | Homo sapiens |
| 32 | MW680969 | Feng,L., Chen,L., Ye,X., Liu,X. | 10784 | 2/2/2018 | China | Homo sapiens |
| 33 | MW680970 | Feng,L., Chen,L., Ye,X., Liu,X. | 10784 | ######## | China | Homo sapiens |
| 34 | MW015936 | Jaimipuk,T., Sachdev,S., Yoksan,S., Thepparit,C., Rajkam,S., Chokephaibulkit,K. | 10807 | ######## | Thailand | Homo sapiens |
| 35 | MN611472 | Shen,Z., Li,X., Lu,Y., Li,J., Xiao,H., Yuan,W., Zhang,Z., Zhou,Y., Feng,Y., Qin,W., Xia,X. | 10807 | 7/1/2019 | China | Homo sapiens |
| 36 | MF996804 | Wongsurawat,T., Athipanyasilp,N., Jenjaroenpun,P., Jun,S.R., Kaewnapan,B., Wassenaar,T.M., Leelahakorn,N., Angkasekwinai,N., Kantakamalakul,W., Ussery,D.W., Sutthent,R., Nookaew,I., Horthongkham,N., Pattama,A. | 10807 | 8/7/2017 | Thailand | Homo sapiens |
| 37 | MG548660 | Wongsurawat,T., Athipanyasilp,N., Jenjaroenpun,P., Jun,S.R., Kaewnapan,B., Wassenaar,T.M., Leelahakorn,N., Angkasekwinai,N., Kantakamalakul,W., Ussery,D.W., Sutthent,R., Nookaew,I., Horthongkham,N., Ussery,D. | 10807 | ######## | Thailand | Homo sapiens |
| 38 | MG548661 | Wongsurawat,T., Athipanyasilp,N., Jenjaroenpun,P., Jun,S.R., Kaewnapan,B., Wassenaar,T.M., Leelahakorn,N., Angkasekwinai,N., Kantakamalakul,W., Ussery,D.W., Sutthent,R., Nookaew,I., Horthongkham,N., Ussery,D | 10807 | ######## | Thailand | Homo sapiens |
| 39 | MG807646 | Wongsurawat,T., Athipanyasilp,N., Jenjaroenpun,P., Jun,S.R., Kaewnapan,B., Wassenaar,T.M., Leelahakorn,N., Angkasekwinai,N., Kantakamalakul,W., Ussery,D.W., Sutthent,R., Nookaew,I., Horthongkham,N., Pattama,A. | 10807 | ######## | Thailand | Homo sapiens |
| 40 | MG807647 | Wongsurawat,T., Athipanyasilp,N., Jenjaroenpun,P., Jun,S.R., Kaewnapan,B., Wassenaar,T.M., Leelahakorn,N., Angkasekwinai,N., Kantakamalakul,W., Ussery,D.W., Sutthent,R., Nookaew,I., Horthongkham,N., Pattama,A. | 10807 | 9/7/2017 | Thailand | Homo sapiens |
| 41 | MH013290 | Wongsurawat,T., Athipanyasilp,N., Jenjaroenpun,P., Jun,S.R., Kaewnapan,B., Wassenaar,T.M., Leelahakorn,N., Angkasekwinai,N., Kantakamalakul,W., Ussery,D.W., Sutthent,R., Nookaew,I., Horthongkham,N. | 10762 | ######## | Thailand | Homo sapiens |
| 42 | MH158236 | Volkova,E., Grinev,A., Gusmao,R., Chancey,C., Rios,M. | 10807 | 2010 | Cambodia | Homo sapiens |
| 43 | MH119185 | Chanama,S., Ruchusatsawat,K., Jampha,W., Wongcharoen,P., Naemkhunthot,S., Okada,P.A., Parnmen,S., Tacharoenmuang,R., Tawatsin,A., Sangkitporn,S., Siriyasatien,P., Tatsumi,M. | 10793 | ######## | Thailand | Homo sapiens |
| 44 | LC369584 | Kato,F., Taniguchi,S., Maeki,T., Tajima,S., Lim,C.K., Saijo,M., Suzuki,R., Takasaki,T. | 10718 | 2017 | Japan:kanagawa, Sagamihara | Homo sapiens |
| 45 | MG674718 | Guo,Y.Q., Fan,F.Y., Bao,L.L., Li,F.D., Lv,Q., Fan,P.H., Jiang,J., Qin,C. | 10807 | ######## | China | Homo sapiens |
| 46 | MG674719 | Guo,Y.Q., Fan,F.Y., Bao,L.L., Li,F.D., Lv,Q., Fan,P.H., Jiang,J., Qin,C. | 10786 | ######## | China | Homo sapiens |
| 47 | KY553111 | Gu,S.H., Song,D.H., Lee,D., Jang,J., Kim,M.Y., Jung,J., Woo,K.I., Kim,M., Seog,W., Oh,H.S., Choi,B.S., Ahn,J.S., Park,Q., Jeong,S.T. | 10795 | 2016-04 | South Korea | Homo sapiens |
| 48 | KX051560 | Melendez,M.C., Klungthong,C., Maljkovic Berry,I., Chusri,S., Buathong,R., Manasatienkij,W., Rutvisuttinunt,W., Chinnawirotpisan,P., Thaisomboonsuk,B., Yoon,I.K., Macareo,L.R., Jarman,R.G., Ellison,D.W. | 10795 | 7/9/2013 | Thailand | Homo sapiens |
| 49 | KX051561 | Melendez,M.C., Klungthong,C., Maljkovic Berry,I., Chusri,S., Buathong,R., Manasatienkij,W., Rutvisuttinunt,W., Chinnawirotpisan,P., Thaisomboonsuk,B., Yoon,I.K., Macareo,L.R., Jarman,R.G., Ellison,D.W. | 10798 | ######## | Thailand | Homo sapiens |
| 50 | KX051562 | Melendez,M.C., Klungthong,C., Maljkovic Berry,I., Chusri,S., Buathong,R., Manasatienkij,W., Rutvisuttinunt,W., Chinnawirotpisan,P., Thaisomboonsuk,B., Yoon,I.K., Macareo,L.R., Jarman,R.G., Ellison,D.W. | 10800 | ######## | Thailand | Homo sapiens |
| 51 | LC219720 | Hashimoto,T., Kutsuna,S., Tajima,S., Nakayama,E., Maeki,T., Taniguchi,S., Lim,C.K., Katanami,Y., Takeshita,N., Hayakawa,K., Kato,Y., Ohmagari,N. | 10696 | ######## | Japan | Homo sapiens |
| 52 | KU761560 | Tong,Y.-G., Wang,G., Tong,Y.-H., Zheng,W., Hu,K.-X., Liu,W., Zhao,T.-Y., Qin,C.-F., Zhang,X.-A., Deng,Y.-Q., Fan,H., Pei,G.-Q., An,X.-P., Mi,Z.-Q., Cheng,S., Gao,B., Cao,W.-C. | 10635 | 2016-02 | China | Homo sapiens |
| 53 | KU761561 | Tong,Y.-G., Wang,G., Tong,Y.-H., Zheng,W., Hu,K.-X., Liu,W., Zhao,T.-Y., Qin,C.-F., Zhang,X.-A., Deng,Y.-Q., Fan,H., Pei,G.-Q., An,X.-P., Mi,Z.-Q., Cheng,S., Gao,B., Cao,W.-C. | 10635 | 2016-02 | China | Homo sapiens |
| 54 | KY272987 | Leelahakorn,N., Horthongkham,N., Sornprasert,S., Athipanyasilp,N., Kantakamalakul,W., Sutthent,R. | 10807 | ######## | Thailand | Homo sapiens |
| 55 | LC191864 | Taira,M., Ogawa,T., Nishijima,H., Yamamoto,K., Hotta,C., Akita,M., Tajima,S., Saijo,M. | 10807 | ######## | Japan:Chiba | Homo sapiens |
| 56 | LC190723 | Ozawa,H., Usuku,S., Nakayama,E., Tajima,S., Kato,K., Yamashita,A., Sekizika,T., Kuroda,M. | 10786 | ######## | Japan:Kanagawa, Yokohama | Homo sapiens |
| 57 | KX694532 | Shabman,R., Puri,V., Dilley,K., Fedorova,N., Shrivastava,S., Amedeo,P., Williams,M., Hu,L., Rashid,S. | 10776 | ######## | Thailand | Homo sapiens |
| 58 | KX694533 | Shabman,R., Puri,V., Dilley,K., Fedorova,N., Shrivastava,S., Amedeo,P., Williams,M., Hu,L., Rashid,S. | 10779 | ######## | Malaysia | Aedes sp. |
| 59 | KX266255 | Bi,Y., Wang,Q., Yang,Y., Liu,Y., Gao,G.F. | 10785 | ######## | China | Homo sapiens |
| 60 | KX601167 | Shabman,R., Puri,V., Dilley,K., Fedorova,N., Shrivastava,S., Amedeo,P., Williams,M., Hu,L., Suthar,M.S. | 10744 | ######## | Malaysia | Aedes sp. |
| 61 | KX377336 | Yun,S.-I., Song,B.-H., Frank,J.C., Julander,J.G., Polejaeva,I.A., Davies,C.J., White,K.L., Lee,Y.-M. | 10807 | 1966-07 | Malaysia | Aedes aegypti |
| 62 | KX253996 | Wu,D., Lin,L., Zhang,Y., Li,J., Liang,M., Li,D. | 10807 | ######## | China | Homo sapiens |
| 63 | KX117076 | Zhang,Y., Sun,Y., Pan,J., Mao,H., Yan,H., Lou,X., Chen,Z., Xia,S. | 10805 | ######## | China | Homo sapiens |
| 64 | KU955593 | Ladner,J.T., Wiley,M.R., Prieto,K., Yasuda,C.Y., Nagle,E., Kasper,M.R., Reyes,D., Vasilakis,N., Heang,V., Weaver,S.C., Haddow,A., Tesh,R.B., Sovann,L., Palacios,G. | 10807 | 2010 | Cambodia | Homo sapiens |
| 65 | KU820899 | Zhang,Y., Sun,Y., Pan,J., Mao,H., Yan,H., Lou,X., Chen,Z., Xia,S. | 10805 | ######## | China | Homo sapiens |
| 66 | KU744693 | Liu,L., Wu,W., Zhao,X., Xiong,Y., Zhang,S., Liu,X., Qu,J., Li,J., Nei,K., Liang,M., Shu,Y., Hu,G., Ma,X., Li,D., Nie,K. | 10676 | 2/6/2016 | China | Homo sapiens |
| 67 | KU681081 | Ellison,D.W., Ladner,J.T., Buathong,R., Alera,M.T., Wiley,M.R., Hermann,L., Rutvisuttinunt,W., Klungthong,C., Chinnawirotpisan,P., Manasatienkij,W., Melendrez,M.C., Maljkovic Berry,I., Thaisomboonsuk,B., Ong-Ajchaowlerd,P., Kaneechit,W., Velasco,J.M., Tac-An,I.A., Villa,D., Lago,C.B., Roque,V.G. Jr., Plipat,T., Nisalak,A., Srikiatkhachorn,A., Fernandez,S., Yoon,I.K., Haddow,A.D., Palacios,G.F., Jarman,R.G., Macareo,L.R., Melendez,M.C., Maljkovicberry,I., Ong-ajchaowlerd,P., Akrasewi,P. | 10807 | ######## | Thailand | Homo sapiens |
| 68 | KU681082 | Ellison,D.W., Ladner,J.T., Buathong,R., Alera,M.T., Wiley,M.R., Hermann,L., Rutvisuttinunt,W., Klungthong,C., Chinnawirotpisan,P., Manasatienkij,W., Melendrez,M.C., Maljkovic Berry,I., Thaisomboonsuk,B., Ong-Ajchaowlerd,P., Kaneechit,W., Velasco,J.M., Tac-An,I.A., Villa,D., Lago,C.B., Roque,V.G. Jr., Plipat,T., Nisalak,A., Srikiatkhachorn,A., Fernandez,S., Yoon,I.K., Haddow,A.D., Palacios,G.F., Jarman,R.G., Macareo,L.R., Melendez,M.C., Maljkovicberry,I., Ong-ajchaowlerd,P., Akrasewi,P. | 10807 | 5/9/2012 | Philippines | Homo sapiens |
| 69 | NC_012532 | Kuno,G., Chang,G.-J.J., Chang,G.J., Tsuchiya,K.R. | 10794 | ######## | Uganda | Simiifor-mes |
